# Social vision expands into auditory cortices of people born deaf – evidence from naturalistic film and encoding models

**DOI:** 10.64898/2026.11.04.763721

**Authors:** Maria Zimmermann, Marcin Szwed, Leyla Isik, Marina Bedny

**Affiliations:** Department of Psychology, University of Warsaw, Warsaw, Poland; Department of Psychological and Brain Sciences, Johns Hopkins University, Baltimore, US; Institute of Psychology, Jagiellonian University, Kraków, Poland

## Abstract

Comparing deaf and hearing adults provides insight into the contribution of experience to function of auditory cortical networks. We used naturalistic fMRI and voxel-wise encoding to measure information processing in auditory networks of deaf (n=24) and hearing (n=22) adults who watched a 35-minute silent animated film. Unlike simpler stimuli, naturalistic films are rich in a broad range of perceptual, affective, and social features, including motion, dynamic social interactions, and Theory of Mind. Machine vision algorithms and human coding were used to densely annotate visual and social film features. In both groups we observed a right-lateralized processing hierarchy with encoding of low-level visual features in early visual areas, social visual features along the superior temporal sulcus (STS) and Theory of Mind in temporoparietal junction. In deaf participants, auditory cortices showed enhanced encoding of a broad range of visual information, from low level motion to Theory of Mind. In higher order auditory networks, social interactions emerging as the strongest predictor of neural activity in deaf people and showed the strongest enhancement in deafness. Auditory networks of deaf adults showed systematic anatomical organization within and across hemispheres. Within the right hemisphere, higher-order auditory regions neighboring the STS preferentially responded to social visual features whereas lower-level visual features were represented broadly across the auditory hierarchy. These results suggest that proximity and connectivity principles govern anatomical expansion of information encoding in sensory loss.

## Introduction

In people born deaf, ‘auditory’ cortices show enhanced responses during visual tasks, including visual motion perception, temporal sequence processing and visual working memory (Bavelier et al., 2001; Bavelier & Neville, 2002; Benetti et al., 2021; Bola et al., 2017; Cardin et al., 2018; Fine et al., 2005; Finney et al., 2001; Manini et al., 2022; Retter et al., 2019; Shiell et al., 2016; Zimmermann et al., 2021). Despite long-standing evidence for such ‘cross-modal’ plasticity, the visual features that are encoded by auditory networks in deafness and the principles that govern experience-based plasticity remain debated.

One challenge is that mechanisms of cortical plasticity in humans are still poorly understood and as a result the space of possible hypotheses about the function of auditory cortex in deafness is large. Classic experimental approaches test one or two hypothesized cognitive functions at a time, leaving many candidates untested. We use voxel-wise encoding models to broadly characterize responses in auditory networks of deaf and hearing adults viewing a silent naturalistic film (Naselaris et al., 2011, 2015; Vu et al., 2011). Naturalistic movies engage many levels of processing simultaneously, from basic perceptual properties to representations of actions, social interactions, mental states, and affect (Im et al., 2025; Lee Masson & Isik, 2021; Redcay & Moraczewski, 2020). This allows us to evaluate a range of candidate cognitive features within the same study and ask which of them are represented in auditory cortices in deafness and how they are anatomically arranged with respect to each other.

Naturalistic films provide an opportunity to examine dynamic social information that is often missed by simpler experimental stimuli. Face and body movements convey information about others’ actions, intentions, and interactions and engage a ‘lateral visual stream’ in superior temporal cortex (Isik et al., 2017; Lee Masson et al., 2024; Lee Masson & Isik, 2021; McMahon et al., 2023; Papeo & Abassi, 2019; Pitcher et al., 2011; Pitcher & Ungerleider, 2021; Wurm & Caramazza, 2019). For people born Deaf, face and body movements play an important role in social communication, most notably for (sign) language recognition (Padden, 1988; Stokoe, 1980). Some evidence suggests that Deaf individuals show enhancements in aspects of face and action perception and enhanced neural responses that extend beyond the language processing context (Benetti et al., 2017; Corina et al., 2007, 2013; Quandt et al., 2021; Simon et al., 2020). Naturalistic films and encoding models provide an opportunity to test whether representations of dynamic social vision expand into auditory cortices in people born deaf.

In the current study, twenty-four deaf and twenty-two hearing participants watched a 35-minute silent animated film (*The Triplets of Belleville*) during fMRI. We used an encoding approach to characterize the visual information represented across auditory cortices. Using a combination of computational vision models and human annotations, we quantified movie features spanning multiple levels of visual and social processing, including low-level perceptual properties (motion energy and color), the presence of faces and people, social interaction, affective content (arousal and valence), and Theory of Mind (ToM). We then tested which visual and social features were represented in auditory cortices, whether their encoding was enhanced in deaf compared with hearing participants, and whether these effects differed across subdivisions of the auditory hierarchy and between hemispheres.

## Results

We characterized the movie’s low-level visual, social, and affective features. Motion energy and HSV were extracted automatically, while independent raters annotated 4-s segments for social interactions, valence, arousal, faces, and number of people. Theory of Mind (ToM) was annotated over longer event-based segments, reflecting characters’ reasoning about others’ thoughts, emotions, or goals. Annotation procedures were adapted from previous naturalistic fMRI studies (Baldassano et al., 2017; Lee Masson & Isik, 2021; see Methods for details). Given correlations among naturalistic stimulus features, we also assessed their interrelationships (Fig. 1B). To relate these stimulus dimensions to neural activity, we trained voxel-wise encoding models using multi-kernel ridge regression. Each stimulus feature was modeled as a separate feature space, allowing us to estimate its contribution to BOLD responses while accounting for the other features in the movie (Fig. 1A).

**Figure 1.**
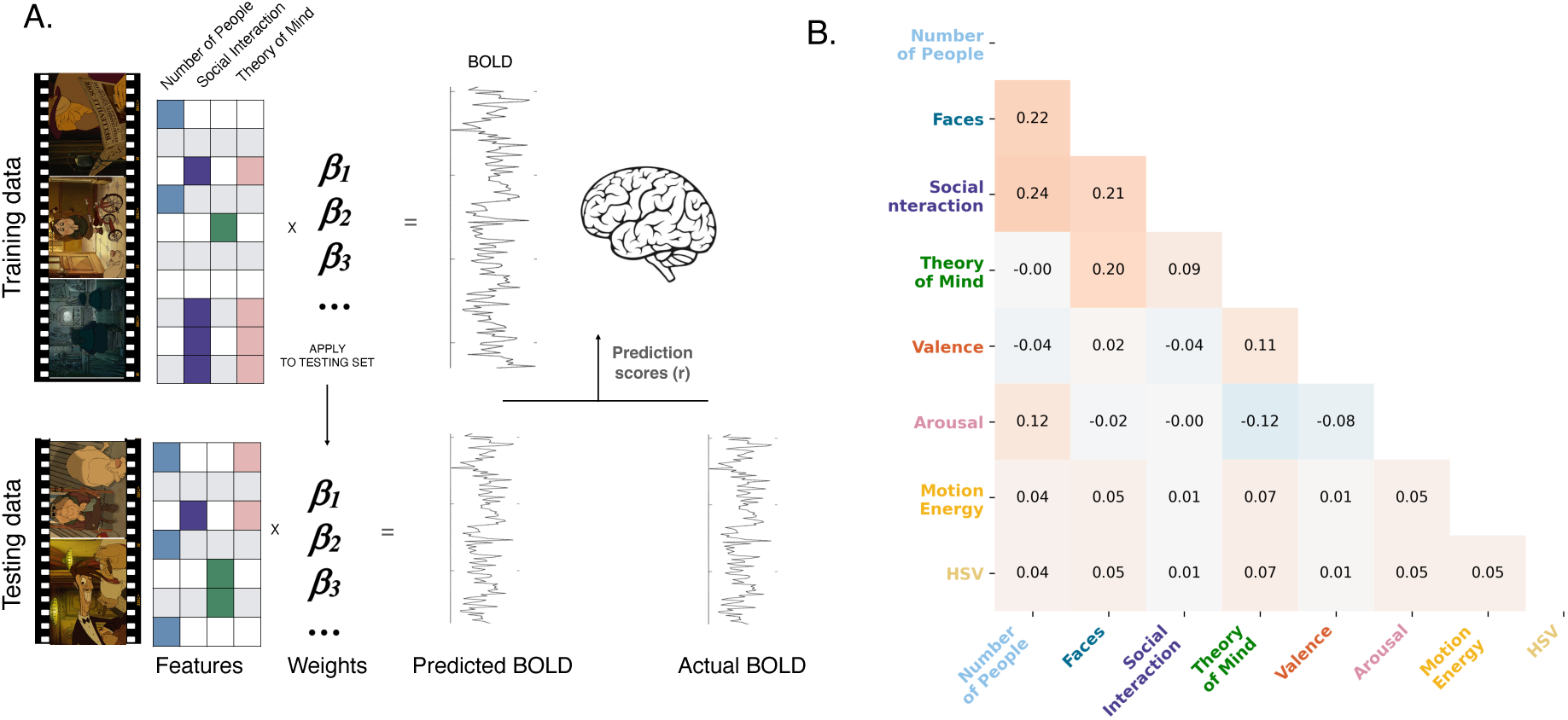
A. Encoding model. Schematic illustration of the voxelwise encoding model used to predict neural responses from stimulus features. B. Between-feature correlations. Correlation matrix showing pairwise relationships among the extracted stimulus features used in the model.

### Enhanced sensitivity to content of silent film in auditory networks of deaf people: full model

Auditory areas of deaf people showed enhanced sensitivity to cognitive content of silent animated film. In the deaf group, the full model, which included all above features, predicted activity in higher-level auditory regions of the superior temporal gyrus (STG) and Sylvian fissure, particularly in the right hemisphere, and posteriorly into the temporo-parietal junction (TPJ).

When groups were compared directly, voxels with significantly higher encoding in the deaf than hearing individuals were located primarily in the lateral temporal cortices, including the STS with extensions into Superior Temporal Gyrus (STG) associated with higher-order auditory processing in hearing individuals, as well as the temporo-parietal junction (TPJ) in the right hemisphere a region commonly linked to theory-of-mind processing.

Anatomically, enhanced responses in the deaf group were adjacent to shared responses along the STS in hearing and deaf people. The full model significantly predicted voxel-wise responses in both hearing and deaf participants along the right superior temporal sulcus (STS) and within visual cortices.

Both the within group effects and between group differences were right-lateralized, with left-hemisphere effects confined to small regions in posterior temporal cortex and TPJ. No voxels showed greater encoding in hearing than deaf individuals.

**Figure 2.**
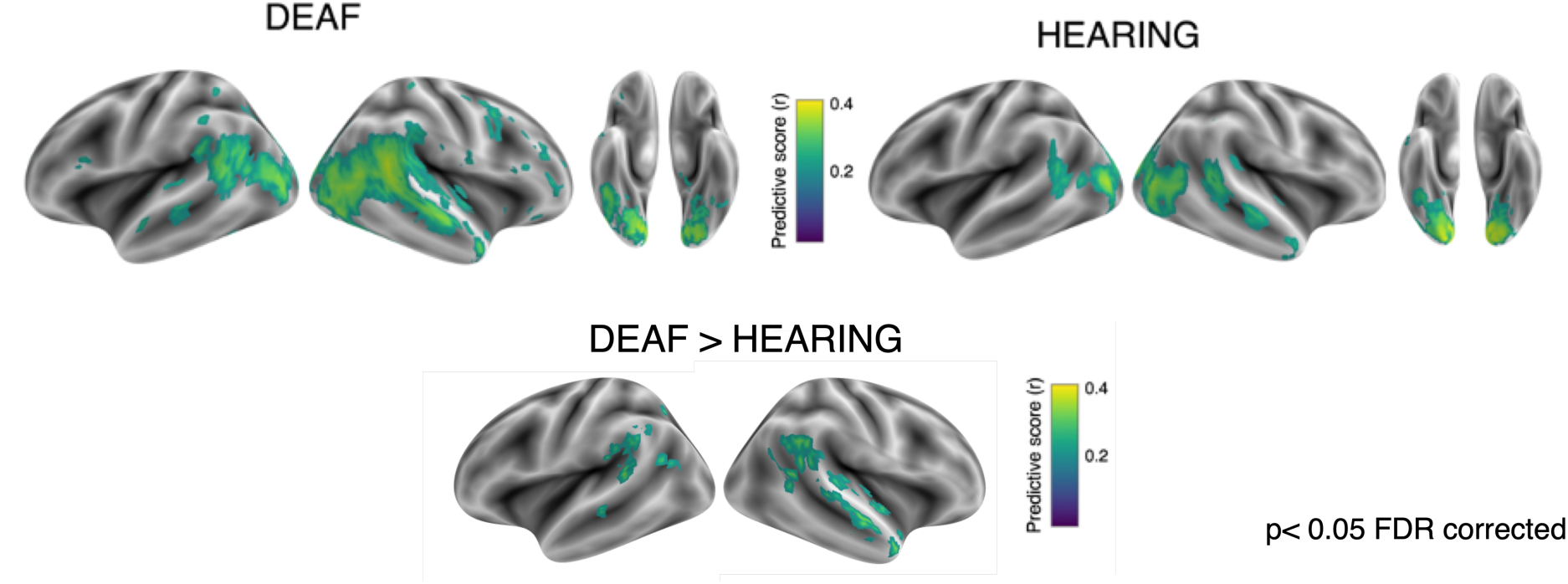
Predictive performance (*r*) of the full models combining all feature types (social, affective, and perceptual), shown separately for each group. Statistical significance was determined using sign-flip permutation, p < 0.05, FDR corrected. The group difference was calculated using a permutation test with group-label shuffling, restricted to voxels with significant predictive scores in either group model: Deaf or Hearing (p < 0.01 uncorrected). Significance was assessed at p < 0.05, FDR-corrected.

### Social features primarily drive enhanced encoding in deaf participant

To probe the cognitive content of enhanced neural predictivity in the deaf group, we applied a winner-takes-all feature analysis to voxels that showed significantly higher predictive scores in the deaf than hearing individuals (Fig. 3). Social interaction emerged as the dominant predictor in the majority of voxels (%) with stronger responses in the deaf group, particularly along the right STS. In addition, posterior portions of the STS and TPJ were most strongly driven by the Theory of Mind feature. Regions in occipitotemporal and lateral occipital cortex showed stronger responses to number of people in the scene and motion energy. This pattern broadly follows the typical processing hierarchy in the lateral occipital and temporal cortices, from low level sensory (e.g., motion) to abstract social (e.g., ToM) features.

**Figure 3.**
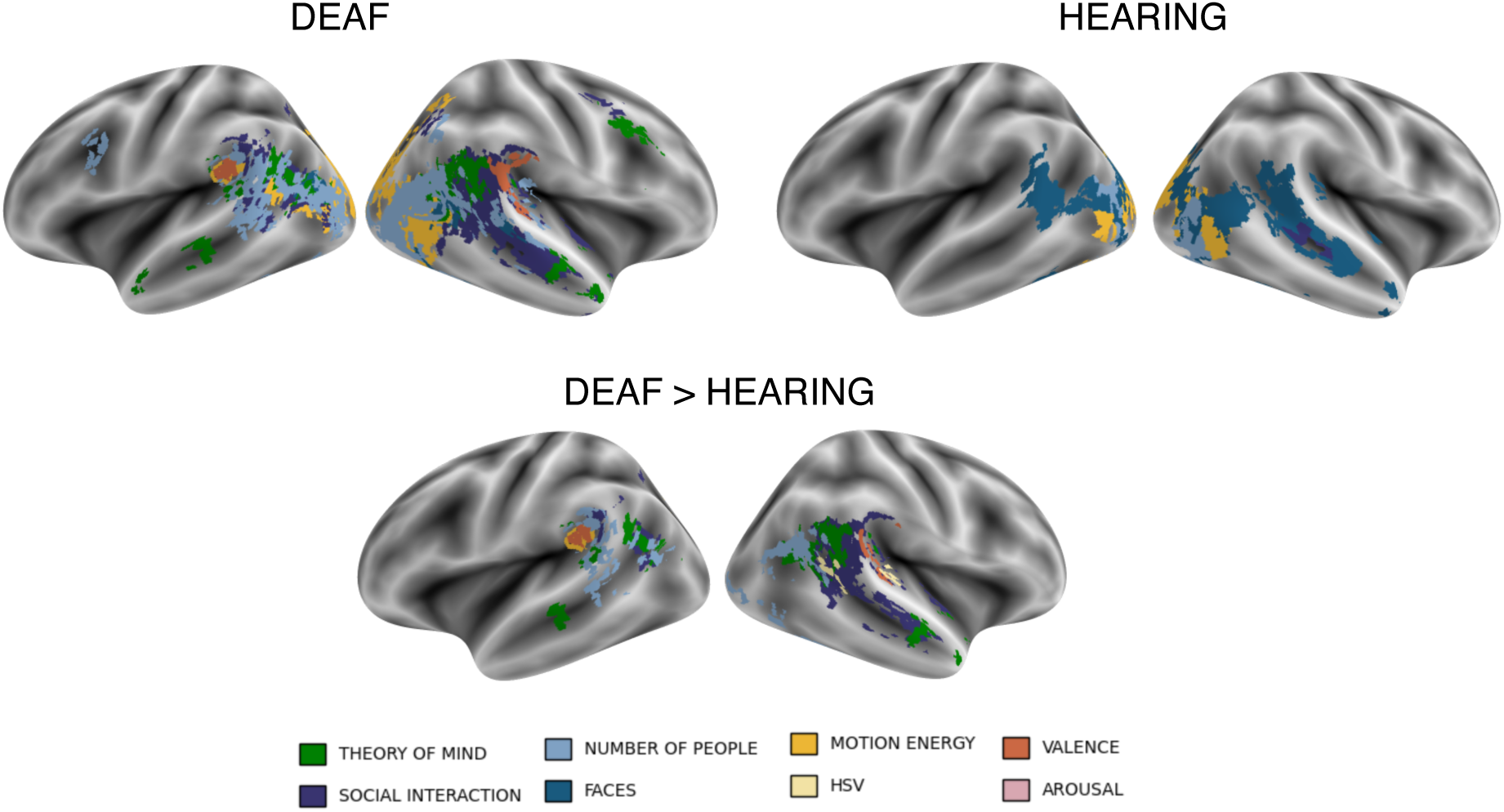
A. Voxels with significant predictive scores in the deaf and hearing groups, shown separately and colored according to winner-takes-all maps indicating the top predictive feature (visual, social, or affective). B. Voxels with significantly higher predictive scores in deaf than in hearing individuals (*p* < 0.001, uncorrected; >10 contiguous voxels), colored according to winner-takes-all maps indicating the top predictive feature in deaf individuals (visual, social, or affective).

Enhanced feature representations in deaf participants were spatially adjacent to corresponding representations observed in hearing individuals (Supplementary Fig. S2), suggesting an expansion into neighboring cortical territory.

Variance partitioning further revealed that three of the social features: social interaction, Theory of Mind, and number of people each uniquely explained variance within different subsets of voxels with enhanced encoding in the deaf group (Fig. S1), demonstrating that these features make independent contributions to the observed responses.

In a direct group comparison in all voxels showing significant responses in either group, social interaction was the only feature with enhanced encoding in the deaf group, centered in the right Superior Temporal Gyrus (Fig. S3.)

### Auditory cortices encode social visual and perceptual features in deaf individuals

We used anatomically defined regions of interest (ROIs) to test whether silent-film features are encoded across auditory and auditory-related temporal cortices in people born deaf, and whether this encoding varies along the processing hierarchy from primary auditory cortex (Te1), through mid-level auditory cortex (Te2–Te3), to multimodal, auditory-related in the upper bank of the mid-STS (STS1/Te4, Belin et al., 2000; Gulban et al., 2020; Nourski et al., 2021; Zachlod et al., 2020, 2022). We refer to this latter ROI (STS1/Te4) as mid-STS throughout. It covers a cytoarchitectonically defined region of the upper STS bank and was included as a higher-order, multimodal extension of the auditory processing hierarchy beyond auditory cortex proper.

Hearing participants showed significant encoding of only a few visual features and only in higher-order auditory and polymodal regions. No feature showed significant encoding in the A1 Referred to as Te1 above (all p’s> 0.05, sign flip permutation test), mid-auditory cortex encoded faces and valence whereas mid-STS encoded faces, valence, and social interaction (p<0.05, permutation test, Fig. 4). This pattern is consistent with increasing integration of visual information along the auditory processing hierarchy.

**Figure 4.**
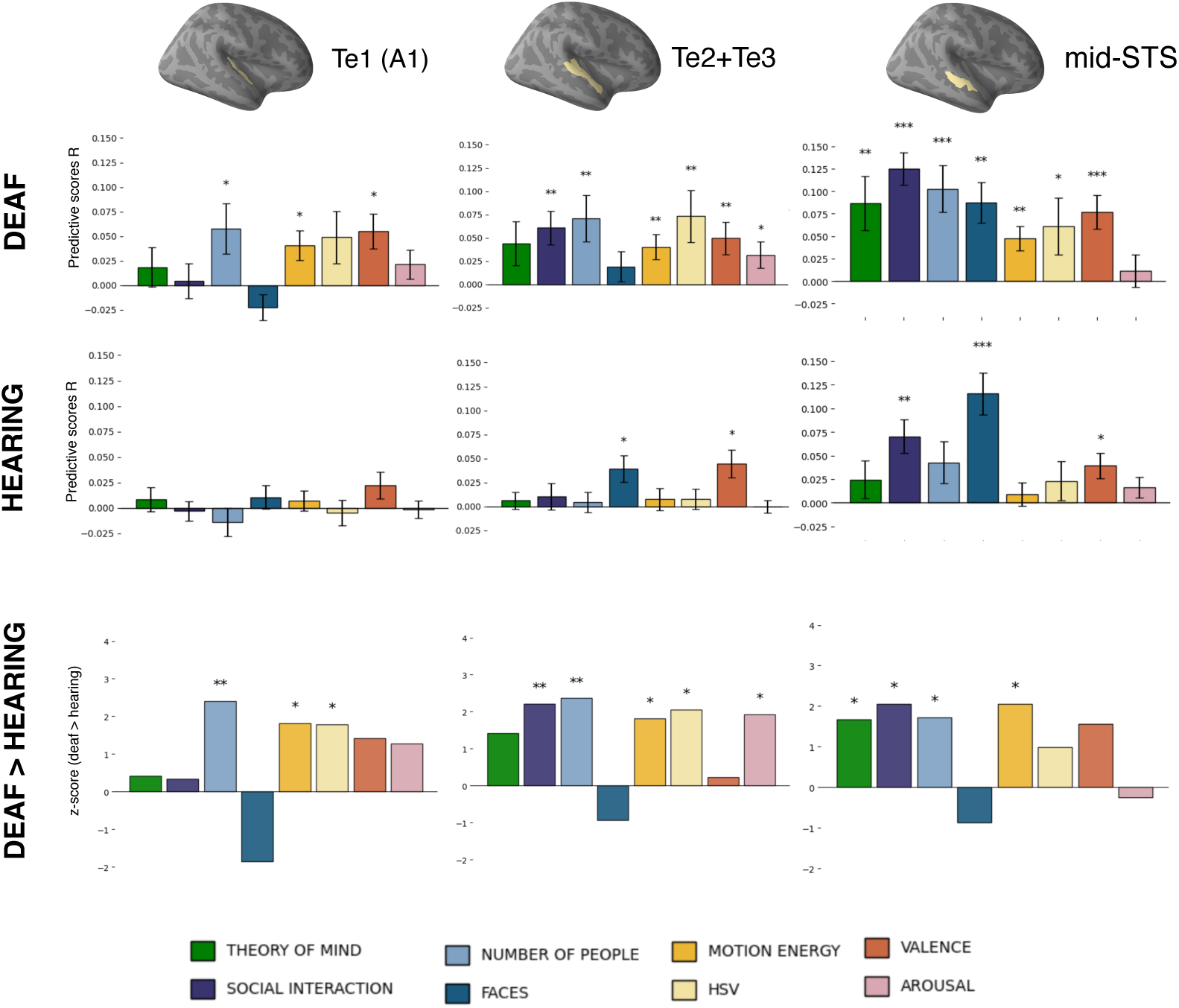
Predictive scores for the deaf and hearing groups for each feature within right-hemisphere auditory ROIs. Within-group significance was assessed using a permutation sign-flip test (*p* < 0.05, \**p* < 0.005, \*\**p* < 0.001), with FDR correction. Between-group differences were assessed using a group-switching permutation test (*p* < 0.05, \**p* < 0.005, \*\**p* < 0.001), with FDR correction.

In contrast, deaf participants showed robust encoding of many silent-film features throughout the auditory hierarchy. Low-level features i.e., motion energy and HSV as well as valence significantly predicted neural responses across all levels of the auditory hierarchy, from A1 to mid-STS, indicating relatively uniform encoding (all feature and ROI p’s < .05, permutation tests, no significant effect of ROI for these features: motion energy, *F*(2, 46) = 0.09, *p* = .917, arousal, *F*(2, 46) = 1.57, *p* = .219, valence, *F*(2, 46) = 1.28, *p* = .287, HSV, *F*(2, 46) = 1.19, *p* = .315.)

Encoding of social features increased along the processing hierarchy in the deaf group in the right hemisphere. We found a significant effect of ROI for social interaction (*F*(2,46) = 28.44, *p* < 0.001) and theory of mind (*F*(2,46) = 4.43, *p* = 0.017) features, as well as faces (*F*(2,46) = 11.58, *p* < 0.001) and a marginal effect for number of people: (*F*(2, 46) = 2.92, *p* = .061), reflecting stronger encoding in higher-order regions. Social interaction showed significant encoding in higher-level auditory cortices and mid-STS and the theory of mind feature was significant only at the highest level of the hierarchy (mid-STS) (p=0.005, permutation test) (Fig.4).

In the left hemisphere of deaf individuals, some features showed significant encoding at each of the three levels of the auditory hierarchy (A1, mid-auditory cortex, and mid-STS; see Fig. S1 for details). Unlike in the right hemisphere, however, the effect of ROI was not significant for any feature in the left hemisphere, providing no evidence for hierarchical specialization.

Notably, face-presence in the scene was the only feature that did not show any encoding enhancement in deaf participants, compared to hearing (for every ROI p>0.05 permutation test). Instead, these features tended to be more strongly represented in hearing individuals (but not significantly) (Fig. 4). Both face and valence significantly predicted responses in higher auditory and polymodal STS cortices in hearing participants. This result suggests that deafness-enhanced responses to social interaction and theory of mind are not merely driven by presence of the face.

## Discussion

During naturalistic film viewing, visual information is represented throughout the auditory cortices of people born deaf, with the strongest responses to dynamic social visual features, particularly social interactions. The superior temporal sulcus represents dynamic social visual information, including body movements, facial movements, and interactions among agents, in both hearing and deaf individuals, consistent with its established role in social perception (Isik et al., 2017; Papeo & Abassi, 2019; Walbrin et al., 2018). In deaf individuals, these representations are enhanced in the STS and extend superiorly into the auditory cortices. Within auditory networks of the right hemisphere, higher-level social visual features, particularly social interactions, are preferentially encoded in higher-order auditory regions adjacent to the right STS, whereas lower-level visual features (e.g., motion energy) are represented broadly throughout the auditory hierarchy.

### Enhanced encoding of visual social interactions in higher-order auditory networks of deaf people

Visual perception of social interactions was the strongest driver of activity in auditory networks of people born deaf. Social interaction predicted activity in large swaths of auditory cortex, particularly in right higher-order auditory regions of deaf individuals. In the animated film, this feature captures scenes in which two or more people engaged in a joint action rather than acting independently. Neural selectivity for social interaction has been consistently linked to regions in the right superior temporal sulcus (STS) in hearing people (Isik et al., 2017; Lee Masson et al., 2024; McMahon et al., 2023). For instance, the STS is more strongly engaged when observing people actively interacting, such as conversing, sharing a meal, or playing sports together, than when viewing non-interactive actions and patterns of activity within the STS distinguish between social interactions (e.g., heling vs. hindering)(Isik et al., 2017; Lee Masson et al., 2024; Papeo & Abassi, 2019; Walbrin et al., 2018). Here, we find that this selectivity is enhanced in deafness and extends superiorly into higher-order auditory regions.

Consistent with previous reports, we found enhanced sensitivity to visual motion energy throughout the auditory networks of deaf people, including in primary auditory (Te1), mid-auditory (Te2-Te3) and polymodal mid-STS regions bilaterally (Bavelier et al., 2001; Benetti et al., 2021; Retter et al., 2019; Shiell et al., 2016). Previous studies using point-light displays have also shown auditory-cortical sensitivity to biological motion in deaf individuals (Simon et al., 2020). Here, we observed sensitivity to the number of people present in the movie scene, a feature that may reflect the presence of moving bodies. However, even when visual motion and the presence of moving bodies were accounted for, social interactions explained unique variance in the right higher-order auditory cortices and adjacent STS beyond – suggesting additional sensitivity to this feature.

Likewise, the enhanced responses for social interaction were not explained by the mere presence of faces. Although dynamic faces elicited robust responses in the right higher-order auditory cortex in both hearing and deaf participants, face responses were not enhanced in deafness. At first glance this observation contrasts with previous findings that shows enhanced responses to faces in deaf individuals in a voice-selective STS region of the auditory cortex (temporal voice area) (Belin, 2006; Belin et al., 2000; Benetti et al., 2017). However, unlike the current study, this prior work presented hearing and deaf people with static images of faces and examined sensitivity to face identity (Benetti et al., 2017). In hearing people STS responses to static faces and static face features (e.g., identity) only weakly as compared to the ventral stream which is well activated by static faces (Bernstein & Yovel, 2015; Gobbini & Haxby, 2007; Haxby et al., 2000; Pitcher et al., 2011). Instead, STS is particularly sensitive to dynamic and changeable facial information, including facial motion and expression (Deen et al., 2015, 2020; Fox et al., 2009; Pitcher et al., 2011; Puce et al., 1998, 2003; Schultz et al., 2013). For example, posterior STS responds strongly to facial movements in hearing people, including movements conveying visual speech (Deen et al., 2020; Puce et al., 2003) and silent speechreading can additionally recruit neighboring superior temporal and auditory cortices (Calvert et al., 1997; Capek et al., 2008). Here we find that dynamic, communicative visual faces recruit lateral temporal cortices robustly and equally in both hearing and deaf individuals. By contrast, responses to visual social interactions are enhanced in deafness.

An open question is why social interaction features are enhanced in deaf people and whether these enhancements are associated with behavioral changes. Visual social interactions are represented in the right STS, and in people born deaf, this functional specialization may extend into neighboring higher-order auditory cortices. Recruiting additional neural resources for processing visual social information may also be behaviorally relevant: Social interactions convey rapidly changing communicative information through faces, bodies, gaze, and gestures, cues that are important for social perception generally but may acquire particular relevance in deaf individuals, for whom access to social communication relies more strongly on the visual modality.

### Functional specialization within auditory networks of people born deaf

In the right hemisphere, we found a functional dissociation between lower- and higher-order auditory regions in congenitally deaf individuals. Higher-level features, including social interactions and Theory of Mind, were represented preferentially in higher-order auditory regions adjacent to STS and TPJ areas that represent these high-level features in deaf and hearing people alike. By contrast, lower-level visual features, such as motion energy, were represented broadly throughout the auditory cortex, including A1. Functional differences across the auditory hierarchy in deafness have also been observed in the temporal dynamics of visual processing, particularly in the right hemisphere: early auditory regions in the right hemisphere exhibit shorter temporal integration windows, consistent with lower-level perceptual processing, whereas higher-order auditory regions integrate information over longer timescales, supporting more complex representations (Zimmermann et al., 2024). A number of previous studies with deaf participants have also reported cross-modal visual responses in higher-order auditory regions, but not primary auditory cortex (Benetti et al., 2021; Bola et al., 2017; Cardin et al., 2018; Shiell et al., 2016). These results suggest that different regions within the auditory hierarchy of people born deaf have partly distinct response properties.

The functional dissociation we observed across the auditory hierarchy even in the absence of auditory experience is consistent with known differences in anatomical connectivity across auditory regions that are present at birth. In non-human primates, the primary auditory cortex (A1) is connected predominantly within the auditory system, whereas surrounding auditory regions have broader connections with multisensory and association cortices (Cammoun et al., 2015; Kaas & Hackett, 2000). In this respect i.e., anatomical isolation from non-auditory regions, A1 stands out even among other primary sensory areas, including V1 and S1/M1, which are more broadly connected (Kaas & Hackett, 2000). Human studies suggest a similar organization, with higher-order auditory regions showing stronger connectivity with areas such as the superior temporal sulcus (STS), temporoparietal junction (TPJ), and anterior temporal cortex (Rauschecker & Scott, 2009; Rolls et al., 2023). These distinct connectivity profiles may contribute to different functional specializations across the auditory hierarchy in deaf and hearing people.

We also observed functional differences across hemispheres. Prior studies find that dynamic social visual perception functions tend to be right lateralized (Isik et al., 2017; Pitcher & Ungerleider, 2021). In the current study our encoding models explained substantially more variance in the right than in the left temporal cortex, in both hearing and deaf groups. Likewise, enhanced encoding in deaf individuals was more anatomically extensive in the right hemisphere. This hemispheric asymmetry is also consistent with evidence from prior task-based studies. Stronger recruitment of the left auditory cortex in deafness has been observed during verbal and working-memory tasks (Manini et al., 2022; Twomey et al., 2017), whereas visual perceptual tasks more consistently recruit the right auditory cortex in people born deaf (Bola et al., 2017; Shiell et al., 2016; Zimmermann et al., 2021). Sign language stimuli robustly engage bilateral temporal networks in people born deaf, including the left hemisphere (Emmorey, 2021; Emmorey et al., 2002; MacSweeney, 2002; Terhune-Cotter & Emmorey, 2025; Trettenbrein et al., 2024; Zimmermann et al., 2026).

In both deaf and hearing individuals, the right STS preferentially processes nonverbal social visual information, whereas the left STS is more strongly involved semantic and linguistic processing (Sueoka et al., 2024). In people born deaf, these functions may extend into adjacent auditory cortex in the same hemisphere through stronger intra than interhemispheric connections. As a result, crossmodal plasticity may recapitulate hemispheric specialization present in adjacent cortical networks.

## Conclusion

Auditory cortices in people born deaf encode visual information during viewing of naturalistic films, with particularly strong responses to social interactions. Lower-level visual features, such as motion, are represented broadly across the auditory hierarchy, whereas higher-level social features, including social interactions, are preferentially encoded in higher-order auditory and polymodal regions, with strongest responses in the right hemisphere. This pattern of specialization suggests that anatomical proximity and connectivity with polymodal cortical regions that process high-order information in the visual domain may contribute to auditory cortex specialization in the absence of auditory experience.

## METHOD

### Participants

Twenty-four deaf participants and twenty-two hearing participants watched a 35-minute silent animated film (*The Triplets of Belleville*) in the fMRI scanner. The film contained no language, and the soundtrack was removed. 20 of Deaf participants and 22 hearing participated in the previous study (for details see (Zimmermann et al., 2024). Hearing participants were matched by sex and age to the deaf participants (mean). Inclusion criteria for deaf participants included congenital or early deafness, severe level of deafness (above 90dB threshold in both ears), and lack of a cochlear implant. For detailed information on deafness etiology and language experience, see Table1. Instructions in Polish Sign Language (PJM) for deaf participants (by use of a sign language interpreter). Written informed consent was signed by all participants.

### Data Acquisition

MRI structural and functional data of the whole brain were collected on a 3 Tesla Siemens MAGNETOM Tim Trio scanner. The T1-weighted structural images were collected in 176 axial slices with 1 mm isotropic voxels using a magnetization-prepared rapid gradient echo (MP RAGE). Functional images were collected using a gradient echo planar imaging (EPI) sequence (60 sequential ascending axial slices, repetition time (TR) 1.4 seconds, echo time (TE) 30ms, flip angle 70°, field of view (FOV), matrix 76 x 70, voxel size 2.5 x 2.5 x 2.5 mm, PE direction L/R, multiband MB=3). Data analyses were performed using fmriprep (freesurfer), python (v. 3.7).

### Data preprocessing

We performed minimal preprocessing using *fmriprep* (Esteban et al., 2019) the same as reported in Zimmermann, et al. 2024. Anatomical preprocessing: T1-weighted images were corrected for intensity non-uniformity (N4BiasFieldCorrection), skull-stripped (antsBrainExtraction, OASIS30ANTs), segmented (FSL FAST), and normalized to MNI152NLin2009cAsym (ANTs). BOLD reference was co-registered to T1w (FSL FLIRT, 9 DOF). BOLD time series were corrected for head motion, resampled to native and MNI space. Functional preprocessing: Reference volumes were generated, and confounds estimated: FD, DVARS, global signals (CSF, WM, whole-brain), and CompCor components (tCompCor, aCompCor). Motion estimates and global signals were expanded with derivatives and quadratics. Frames with FD > 0.5 mm or DVARS > 1.5 were flagged as outliers. Denoising: Data were smoothed (6 mm FWHM) and denoised with a voxel-wise GLM including 24 motion parameters, outlier regressors, linear/quadratic trends, and CSF signal.

### Movie Annotations

We used a machine vision algorithm to automatically extracted visual motion features for each movie segment using the Adelson and Bergen motion energy model (Adelson & Bergen, 1985), as implemented in the *pymoten* software package (Nunez-Elizalde et al., 2021). Hue – saturation-value (HSV) color components were likewise extracted automatically using an OpenCV algorithm (Bradski & Kaehler, 2000). Specifically, we computed the average HSV per pixel, then averaged across all pixels in each frame, and averaged these values within each timepoint (1.4 seconds).

### Human ratings

Three independent raters labeled social and affective features of the movie. For every 4.2-second segment (3 TRs), they annotated the presence of social interaction, arousal, and valence. Social interaction was defined as two agents acting toward or communicating with one another in a directed manner. Valence and arousal were each rated on a 5-point Likert scale, with valence ranging from very unpleasant (1) to very pleasant (5), and arousal from very calming (1) to very exciting (5). These labels and rating scales were adapted from prior work using movie stimuli in fMRI studies (Lee Masson & Isik, 2021). Additionally, one rater labeled each 4.2 second segment for the presence of faces (yes/no) and categorized the number of people on screen (none, one, two, or more).

Theory of Mind (ToM) content was annotated by three additional raters using longer non-overlapping segments (∼20–30 TRs). For this annotation procedure, the movie was divided into longer segments using a movie segmentation task. The procedure based on (Baldassano et al., 2017) in which raters indicated where the boundaries of events are in the movie and what is the meaning of each segment. To indicate the presence of Theory of Mind, raters indicated whether a segment involved characters reasoning about the mental states such as thoughts, emotions, or goals of others. They additionally explained their choice by indicating what the character is most probably thinking or reasoning about (for example “*Grandma found a hut and thought her grandson must be in trouble”*). All raters watched the entire movie before completing the rating task. Raters received the following instructions and explanations for the rating task:

#### Number of People

Are there people in the scene, and if so, how many? Count all people visible in the scene

#### Faces

Is a face or are faces visible in the scene? Indicate whether at least one face is visible. A dog’s face should also be counted.

#### Social Interaction

Does the video clip show a social interaction between two or more people? A scene with no social interaction may involve one or more people engaging in solitary, non-interacting actions, such as sleeping, walking, or taking a bus. A non-interaction scene may also contain no people. A scene with social interaction includes two or more people interacting with one another, for example talking, having dinner together, hugging, or engaging in a joint action.

#### Valence

How pleasant is the event? Rate the overall emotional tone of the scene. Positive events (e.g., a celebration) are pleasant, while negative, unpleasant events (e.g., an argument or loss) are unpleasant. Events that are neither positive nor negative are neutral.

#### Arousal

How emotionally intense or arousing is the event for you? Rate the emotional intensity of the event, regardless of whether it is positive or negative. An arousing event may be upsetting, joyful, or exciting. A calm event is neutral and does not evoke strong emotions. Theory of Mind (ToM): Does the scene involve a character thinking or reasoning about another character’s mental state, such as their thoughts, emotions, intentions, beliefs, or goals? If yes: Briefly describe what the character thinks, believes, feels, or infers about the other character.

### Inter-rater reliability

was assessed using Spearman’s rho. Annotations showed good consistency across raters, with the lowest reliability for arousal (ρ = 0.48) and valence (ρ = 0.68), and the highest for Theory of Mind (ρ = 0.82) and social interaction (ρ = 0.85).

### Between feature correlation

To examine relationships between visual and social features, we performed representational similarity analysis (RSA) (Kriegeskorte, 2008)). Representational dissimilarity matrices (RDMs) were computed for multidimensional features (motion energy and HSV color) using Pearson correlation distance (1 – Pearson’s *r*) across all pairs of movie segments (TRs). For one-dimensional annotations, dissimilarities were defined as the absolute differences in mean ratings between each pair of segments. To assess feature similarity, we correlated the lower triangles of each pair of feature RDMs. (Fig.1B).

### Encoding Model

To relate brain activity to specific movie features, we trained a voxel-wise encoding model using multi-kernel ridge regression to estimate beta weights linking annotated stimulus features to fMRI BOLD responses. The model was implemented using the *Himalaya* package (Dupré La Tour et al., 2022, 2025), with each feature space (e.g., arousal, valence, faces etc.) treated as a separate kernel. Regularization strength was optimized independently for each feature space and voxel using nested cross-validation with randomized hyperparameter search.

The data were divided into 70 continuous, non-overlapping segments of 20 TRs each (TR = 1.4 seconds). Of these, 7 segments were randomly assigned to the test set, and the remaining 63 segments were used for training. The training set was further divided into 10 folds for parameter selection using leave-one-fold-out cross-validation. Within each outer fold, the regularization parameters for all feature spaces were optimized via randomized search using a 5-fold inner cross-validation loop. To account for the hemodynamic delay, we included five temporally shifted copies of each feature, corresponding to delays of 1 to 5 TRs. These delayed features were concatenated and passed to the model, allowing it to learn voxel-specific temporal response profiles.

### Statistical Analysis

Statistical significance of encoding was assessed using permutation-based tests. To determine whether encoding within each group was significantly greater than chance, we used one-sided sign-flipping permutation tests with 10,000 permutations to generate null distributions. To compare encoding between deaf and hearing participants, null distributions were generated by randomly shuffling group labels and recalculating the difference between group means 10,000 times. In both analyses, the observed effects were compared with the corresponding null distributions to determine statistical significance. For whole-brain analyses, voxel-wise statistical results were corrected for multiple comparisons using the Benjamini– Hochberg false discovery rate (FDR) procedure (Benjamini & Hochberg, 1995).

For ROI analyses, we used mixed ANOVA to test the effects of group and ROI and their interaction on overall encoding. Within the deaf group, repeated-measures ANOVAs were performed separately for each feature to test whether encoding varied across ROIs. The resulting *p* values were corrected for multiple comparisons across features using the Benjamini–Hochberg FDR procedure.

### ROI definition

Regions of interest (ROIs) in the auditory cortex were defined following the approach described in Zimmermann et al. (2024), based on the Julich atlas. Specifically, we identified four auditory regions spanning the cortical auditory processing hierarchy: the primary auditory cortex (Heschl’s gyrus; Te1), higher-order auditory cortex (Te2 and Te3) within the superior temporal gyrus (STG), auditory-related region in the upper bank of the mid-STS (Zachlod et al., 2020). We refer to the latter ROI as mid-STS throughout Together, these regions encompass successive processing stages from primary auditory cortex to to multimodal, auditory-related temporal cortex.

## Data Availability Statement

The data supporting the findings of this study are currently available from the corresponding author upon reasonable request. The associated datasets and analysis materials will be made publicly available through the Open Science Framework (OSF) upon publication of the peer-reviewed article.

## Funding

This study was supported by funding National Science Centre, Poland (grant no. 2024/52/C/HS6/00207 to M.Z. and grant no. 2018/30/A/HS6/00595 to M.S.) M.Z. is supported by the National Science Centre, Poland and Johns Hopkins University; Computations were carried out with the support of the Interdisciplinary Centre for Mathematical and Computational Modelling, University of Warsaw (ICM UW), under the computational allocation awarded to M.Z. (g100-2194).

## Notes

### Competing Interest Statement

The authors have declared no competing interest.

