## Supplementary figures for "Social vision expands into auditory cortices of people born deaf – evidence from naturalistic film and encoding models"

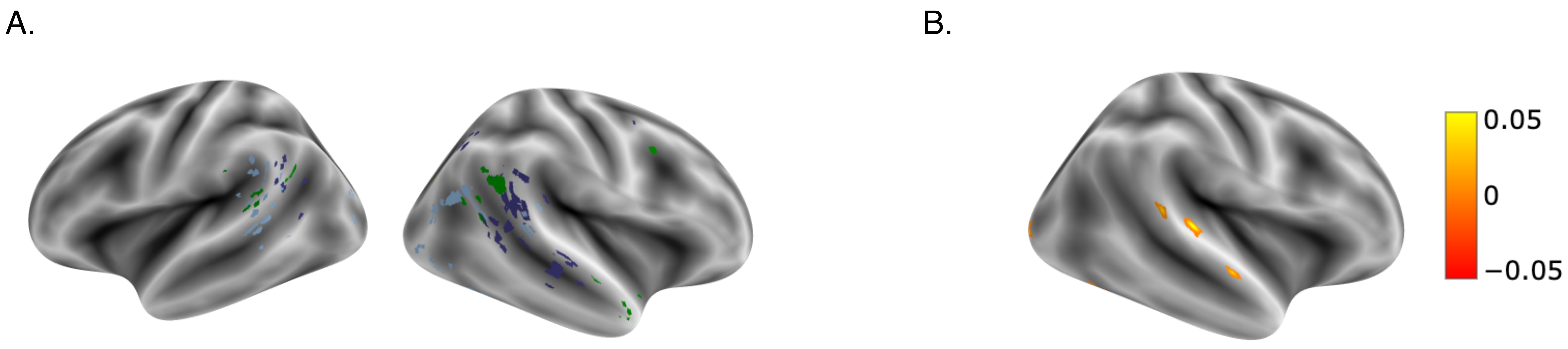


S1. A. Voxels uniquely explaining variance in the deaf group (colored by the corresponding feature). B. Predictive scores DEAF > HEARING for social interaction feature, p<0.05, FDR corrected.

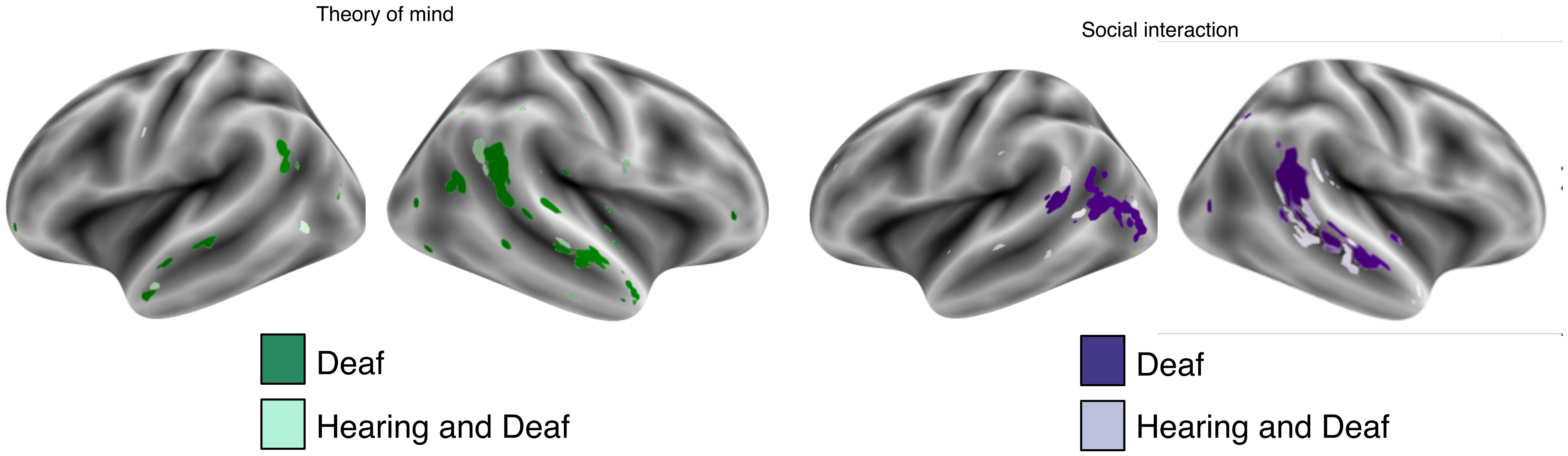


S2. Voxels showing significant prediction accuracy (Pearson r) in Deaf participants, Hearing participants and group overlap for social features, p<0.001 cluster corrected.

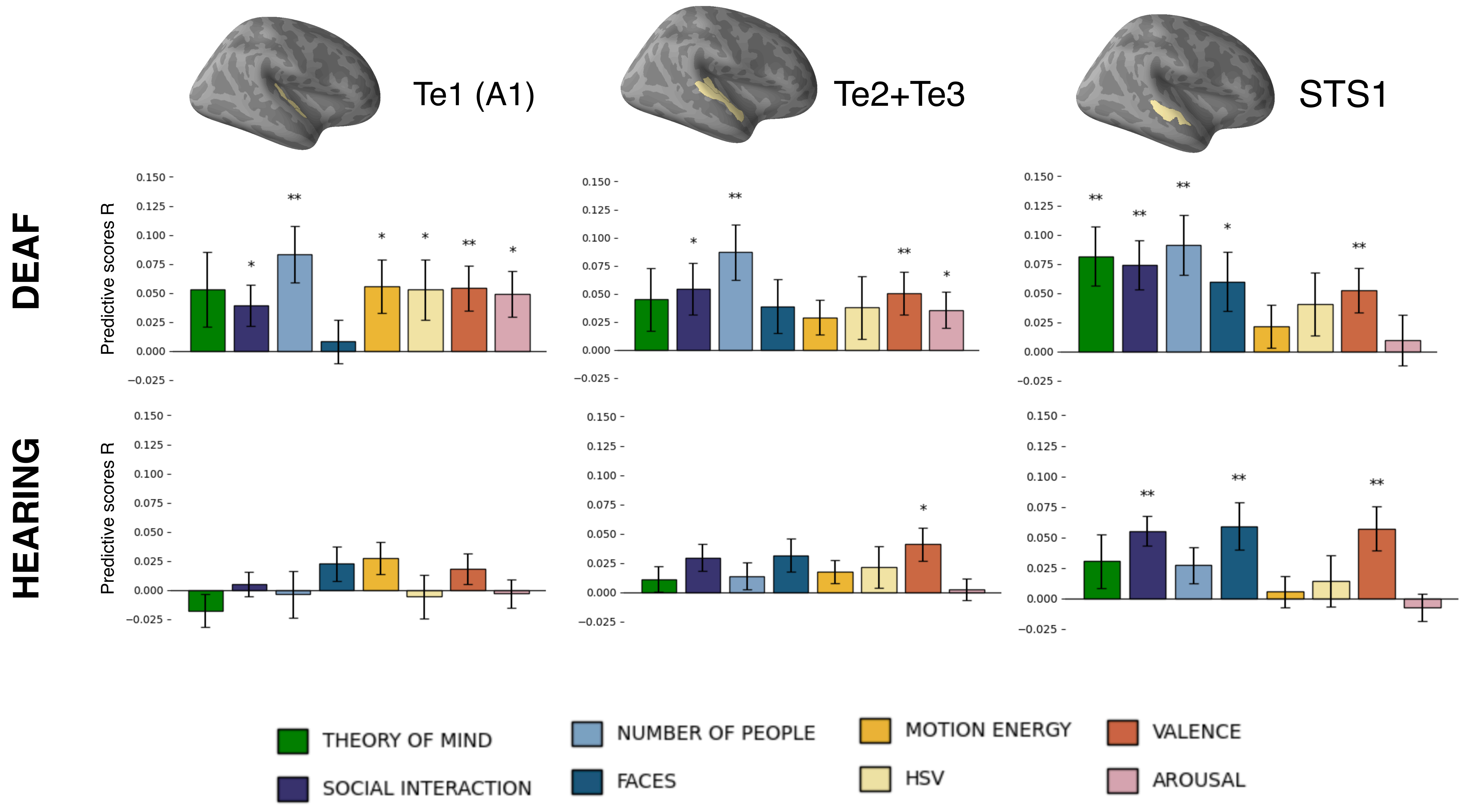


S3. Predictive scores for deaf and hearing groups for each feature within **left-hemisphere auditory ROIs.** Significance computed using permutation test ($p < 0.05$, ** $p < 0.005$, *** $p < 0.001$), FDR corrected, error bars represent SEM.

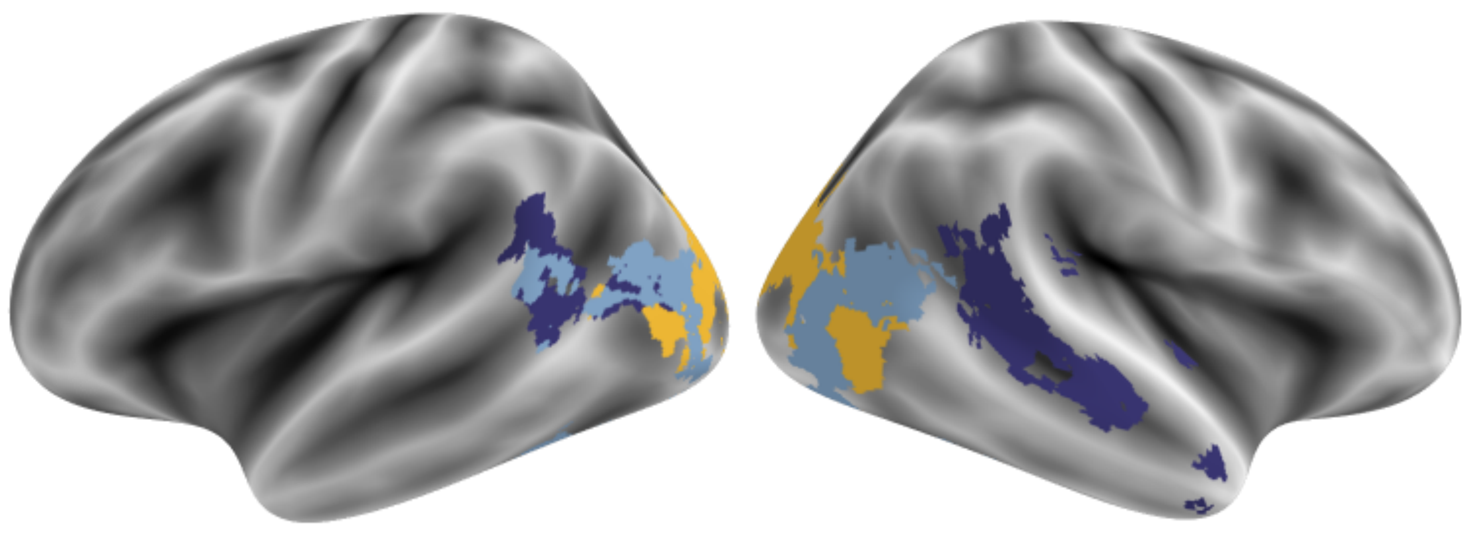

S4. Voxels with significant predictive scores in the hearing group, colored according to a second-winner-takes-all map indicating the feature (visual, social, or affective) with the second-highest predictive score after excluding the top-ranked feature.

Table1.

| | sex | age | education | level of deafness | onset of deafness (diagnosis of hearing loss at the current level) | hearing aids | deaf mother | deaf father | PJM native | Polish Sign Language fluency |
|---|---|---|---|---|---|---|---|---|---|---|
| 1 | F | 33 | higher | > 120 dB | congenital | No | No | No | No | fluent |
| 2 | M | 37 | higher | 90- 119 dB | congenital | No | No | No | No | fluent |
| 3 | M | 32 | higher | 90- 119 dB | congenital | Yes | No | No | No | fluent |
| 4 | M | 29 | vocational | 90- 119 dB | 5 months | No | No | No | No | fluent |
| 5 | F | 36 | secondary | 90- 119 dB | congenital | No | Nie | Yes | Yes | fluent |
| 6 | M | 23 | secondary | > 120 dB | congenital | No | Yes | Yes | Yes | fluent |
| 7 | M | 27 | higher | 90- 119 dB | congenital | Sometimes | Yes | Yes | Yes | well |
| 8 | M | 34 | higher | 90- 119 dB | congenital | No | Yes | Yes | Yes | fluent |
| 9 | M | 26 | higher | > 120 dB | congenital | Sometimes | Yes | Yes | Yes | fluent |
| 10 | M | 35 | vocational | > 120 dB | congenital | No | Yes | Yes | Yes | fluent |
| 11 | F | 37 | higher | 90- 119 dB | congenital | No | Yes | Yes | Yes | fluent |
| 12 | F | 28 | higher | > 120 dB | congenital | No | Yes | Yes | Yes | fluent |
| 13 | M | 31 | secondary | > 120 dB | 8 months | No | No | No | No | fluent |
| 14 | M | 34 | higher | 90- 119 dB | 6 months | Yes | No | No | No | fluent |
| 15 | M | 39 | secondary | 90- 119 dB | congenital | Yes | No | No | No | well |
| 16 | F | 21 | higher | > 120 dB | congenital | no | Yes | Yes | Yes | fluent |
| 17 | F | 37 | secondary | 90- 119 dB | 8 months | Yes | No | No | No | well |
| 18 | F | 21 | higher | 90- 119 dB | congenital | Yes | Yes | Yes | Yes | fluent |
| 19 | F | 56 | secondary | 90- 119 dB | congenital | No | Yes | Yes | Yes | fluent |
| 20 | F | 34 | secondary | 90- 119 dB | congenital | Yes | Yes | Yes | Yes | fluent |
| 21 | F | 36 | secondary | > 120 dB | congenital | No | Yes | Yes | Yes | fluent |
| 22 | M | 36 | secondary | > 120 dB | congenital | No | Yes | Yes | Yes | fluent |
| 23 | M | 39 | secondary | 90- 119 dB | congenital | No | Yes | Yes | Yes | fluent |
| 24 | F | 36 | secondary | 90- 119 dB | congenital | No | Yes | Yes | Yes | fluent |

Table1. Demographic and Language Background of Deaf Participants
